# Combining ability analysis of *Cucurbita moschata* D. in Cote d’Ivoire and classification of promising lines based on their gca effects

**DOI:** 10.1101/2024.06.16.597849

**Authors:** Badié Arnaud Kouago, Dagou Seka, Kouakou Fulgence Brou, Beket Severin Bonny, Konan Henri Joel Koffi, Koffi Adjoumani, Raoul Sylvère Sie

## Abstract

Twelve hybrids and their four parental lines were evaluated for their general performances, combining abilities, potency ratio and heterosis effects. Taller plants and smaller fruits characterized the parental line *Long*, while the *Soubre* parent showed favorable fruit and seed traits but with a small number of seeds. The line *Zouan-H* exhibited performances that are close to *Soubre* but with a smaller number of fruits per plant, higher number of seeds per fruit and the weight of the fruit is one-half that of the *Soubre* line. The *Oval* parent is characterized by a high number of seeds per fruit, smaller fruits and shorter plants. Significant effects of heterosis were observed in different hybrids for different characters. And the computed values of potency ratio indicated that gene(s) expressions were characterized by partial dominance in many cases and super dominance in some other cases. The variance of the effect of the general combining ability of the female parent was highly significant for all the evaluated traits. The variance of the effect of the general combining ability for the male parent was significant for some characters. The variance due to the effect of the specific combining ability was also highly significant for all the traits evaluated. It showed that the effects of additive genes and non-additive genes contributed to the expressions of the characters. However, the general predictive ratio was closer to 1.00 for all the fruit-related traits implying the strong effects of additive genes in the determination of the fruit-related characters. A classification of the parental lines based on the effects of their general combining ability grouped the *Soubre* lines as promising contributors to fruit yield. The parental lines *Long* and *Oval* formed another group likely on the basis of the small size of their fruits, the small pulps, the smaller number of fruits per plant and the high number of seeds per fruit. *Long* would be a candidate parent for the development of cultivars with longer vegetative growth. The parental line *Zouan-H* formed the third group and it was mostly characterized by its advantageous seed traits.

## Introduction

The world population and particularly the African population have recorded a considerable demographic explosion since 1980 [1]. This strong demographic growth requires a boost in agricultural production in order to feed the continuously increasing world population [2]. Anticipation of a food insecurity, a risk of malnutrition and undernourishment lend to increased research activities in agronomy and the development of food crops with high nutritional values. Crops that were previously neglected and classified as minor crops are now being considered as part of the solution to possible future food shortages, food insecurity, and malnutrition [3].

Species from the Cucurbitaceae family are among the neglected and minor crops. However, the Cucurbitaceae family should be considered as one the most important botanical families for humans [4] due to their richness in nutrients and their abundance. *Cucurbita moschata* D., known as butternut squash, is a member of that family, and it is an important food legume for human consumption. Butternut squash is relatively high in energy and carbohydrates. It is a good source of vitamins and its contents of carotenoid pigments and minerals are particularly high [5]. It has a very high potential to cover the nutritional needs of the population, particularly vulnerable groups with regard to the need for vitamin A [6]. *Cucurbita moschata* D. is one of the most used vegetables in healthy diet due to its nutritional benefits. It is also used in medicine in many developing countries because of the bioactive compounds that can be obtained from its seeds and fruits [7].

Despite its many nutritional and medicinal benefits, butternut squash remains a marginalized crop. In Cote d’Ivoire, the yield of butternut squash is very insignificant, and the crop has totally been ignored in agricultural research programs. Given all the nutritional potential of this species, consideration should be given to this crop in agronomic research in order to improve its productivity, its fruit and seed yields and its other characteristics of nutritional values to humans such as its leaves. The productivity of this species can be increased by improving the genetic architecture across different cultivars or through the selection of high yielding cultivar [8]. For this, knowledge of certain genetic parameters that govern the important agronomic traits of the crop is necessary, and can be obtained through diallel cross design [9] and implementation. Diallel cross design is used in plant and animal genetic research to estimate general combining ability (gca), specific combining ability (sca), potency ratio and heterosis for a population from randomly chosen parental lines [10]. Combining ability analysis helps to identify superior parents to be used in breeding programs or to identify promising cross combinations for cultivar development [11]. Crop breeders typically utilize combining ability analyses to choose parents with high general combining ability and hybrids with high specific combining ability effects. General combining ability is a measure of additive gene activity that relates to the average performance of a genotype in a series of hybrid combinations. Specific combining abilities evaluate the average performance of certain hybrid combinations compared to the parental lines and is the result of dominance, epistatic deviation, and genotype environment interactions. Therefore, both gca and sca effects are important in the selection or development of breeding populations [12]. They are referred to in the selection of parents with the potential to produce hybrids exhibiting greater heterosis. Heterosis is a quantifiable, trait-dependent and environment-specific phenotype, which is the performance of the F1 hybrid exceeding that of the parents [13].

The objectives of this study were to: 1) study the combining abilities of the parental lines and the progenies from the crosses on the fruit, seed traits and plant height; and 2) to classify parental lines based on their gca effects in the *Cucurbita moschata* D. germplasm development and the improvement of cultivars of butternut squash.

## 2. Materials and Methods

### 2.1. Plant material and field experiment

The plant material is composed of the seeds of four inbred lines of butternut squash and their F1 hybrids. They are the parental lines *Oval* (*O*), *Long* (*L*), *Zouan-H* (*Z*), and *Soubre* (*S*), and their F1 hybrids resulting from the complete diallel cross of these four lines. The seeds of these parental lines come from accessions originating from Korhogo, Ferkessédougou, Zouan-Hounien and Soubre, respectively, (Table 1). Accessions of the cultivars *Oval, Long, Zouan-H* and *Soubre* first underwent four successive cycles of self-fertilization to create the respective inbred lines, *O, L, Z, S*, used in a complete diallel cross to obtain the seeds of the different families of F1 hybrids. A total of twelve hybrids and the four inbred lines were used in this study.

**Table 1:** Four accessions of *Cucurbita moschata D*. used in this study and their origins.

| Accession | Name | One-letter name | Origin |
| --- | --- | --- | --- |
| Korho | *Oval* | *O* | Korhogo (North) |
| Ferke | *Long* | *L* | Ferkessédougou (North- East) |
| Zh | *Zouan-H* | *Z* | Zouan-Hounien (West) |
| Soubre | *Soubre* | *S* | Soubré (South-West) |

We conducted the experiment from June to November 2022 in a farming environment on a field of 5673 m^2^ (93 m x 61 m) using a randomized complete block design with three replications. Each block measured 93 m by 18 m (1674 m^2^), and included 16 plots, randomly made of the twelve hybrids and the four parental lines. Each plot had a surface area of 54 m^2^ (9 m x 6 m) with 3 sowing lines and 4 sowing points per sowing line. Only one F1 hybrid or one parent is randomly assigned to a plot. The experimental field had a total of 576 plants. All the other agronomic management practices, such as weeding, hoeing were applied as recommended, and as needed.

### 2.2. Data Collection and Analysis

We collected data at harvest on a sample of 10 plants randomly selected from each of the four parental lines and twelve F1 hybrids in each block. We evaluated the hybrids and the parents based on 12 quantitative characters listed in Table 2. The data collected served to compute the descriptive statistics for those characters. Then, we conducted an analysis of variance with the measured traits serving as the response variables. Blocks and the 16 genotypic strains were the factors. When significant differences were observed, we used the method of Tukey to separate means due to the genotypic effect of the strains at the 5% level of probability. In addition, we determined the effects of the general combining ability (gca) of parent *i* and parent *j* and the specific combining ability (sca) of the cross between two parents *i* and *j* as explained below [9]:

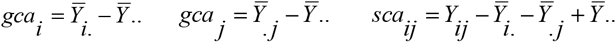

Where *gca*_*i*_ and *gcaj* are the *gca* effects of parent *i* and parent *j*, respectively, and *sca* is the *sca* effect of the cross, 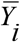 is the mean of all crosses with the same parent *i*; *i*. 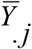 is the mean of all crosses with the same parent *j*; *Y* is the mean of the crosses between parent *i* and parent *j*; and *Y* .. is the mean of all crosses. We assessed the genetic variances of the gca and sca effects with the linear diallel model given below [9]:

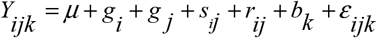

where *Y* is the observed response value of the cross between parents *i* and *j*; μ is the overall mean; *gi* is the general combining ability effect of parent *i*; *g j* is the general combining ability of the parent *j*; *s* is the specific combining ability of the cross between parents *i* and *j*; *r* is the effect of the reciprocal cross between parents *i* and *j*; *b* is the effect of the *k* ^*th*^ block; and *ε*_*ijk*_ is the experimental error.

**Table 2:** Traits of *C. moschata* evaluated in this study, along with their descriptions and unit of measure in parenthesis.

| Abbreviated Name of Traits | Description and (unit of measure) |
| --- | --- |
| PLH | Plant height (cm). |
| NFP | Number of fruits per plant (unit). |
| LOF | Length of fruit (cm). |
| LAF | Width of fruit (cm). |
| DOF | Fruit diameter (cm). |
| WOF | Weight of fruit (g). |
| WTM | Weight of the pulp (g). |
| NOS | Number of seeds per fruit (unit). |
| WFS | Weight of the fresh seeds per fruit (g). |
| W100 | Weight of 100 dry seeds per fruit (g). Obtained after placing the fresh seeds in an oven for 3 days at 70°C. |
| LDS | Length of a dry seed (mm). Average of longest axis of 30 seeds |
| WIS | Width of a dry seed (mm). Average of second longest axis of 30 seeds. |

We determined the relative importance of the gca and sca effects with the general predictive ratio (*gp*r) [14, 15] given as follows:

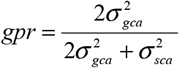

where 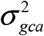 is the variance of the *gca* effect and 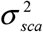 is the variance of the *sca* effect. A *gprsca* closer to unity indicates the hybrid’s performance is a function of the *gca* alone [14]. We estimated the percentage of heterosis with two criteria, the mid-parent heterosis (*mph*) and the better-parent heterosis (*bph*). They are:

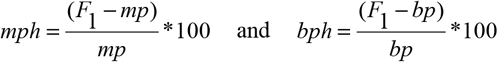

Where *F*_1_ is the mean performance of the *F*_1_ hybrid, *mp* = (*P*_1_ + *P*_2_) / 2 is the average value of the two parents, *P*_1_ and *P*_2_, and *bp* is the value of the better parent. The two statistics, *mph* and *bph*, have a normal distribution [16, 17]. Accordingly, a significance test for a sample of size *n* has been developed with the following test criteria:

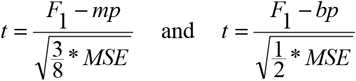

respectively for the mid-parent heterosis and the better-parent heterosis. The statistic *t* has a Student *t*-distribution with (*n* – 1) degrees of freedom and *MSE* is the error variance. The potency ratio (*P*) determined the degree of gene dominance and was computed as follows [18, 19]:

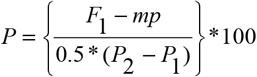

With *P*_2_ representing the average performance of the better parent, and *P*_1_ the average performance of the weaker parent. We should mention that the space of *P* is the set of real numbers. However, *P* = ±1 % indicates complete dominance, − 1% < *P* < 1% means partial dominance, *P* = 0% means absence of dominance and *P* > 1% or *P* < −1% is the indication of super-dominance. The positive or negative sign of *P* defines the direction of dominance with respect to the parents [18]. Randomly selected individuals of each parental line were used to create classes of promising parental lines based on their *gca* effects on plant height and fruit- and seed-yield traits. The statistical software R [20], the packages lmdiallel [21] and ape [22] and the online application iTOL [23] were used for data analysis and graphics.

## 3. Results

### 3.1 Mean Performances of Parents and Hybrids

The means of the plant height, fruit- and seed-yield traits are reported in Table 3. The analysis of variance showed highly significant differences among the parental lines and the hybrid combinations for all the traits studied. Figure 1 shows the diversity of the fruit traits among the parental lines. The method of Tukey for the multiple comparison of means helped to identify significantly different means followed with different letters. It could be seen that the lines identified as *Long* have the highest plant height among the four parental lines. The hybrids involving *Long* as a parent have the highest plant height. The *Soubre* parent is characterized by higher number of fruits that have larger dimensions (length, width, and diameter of the fruit) and are the heaviest. Hybrids produced with *Soubre* as female have the largest number of fruits, the longest, widest and heaviest fruits. In addition, the fruit of the *Soubre* parents have the heaviest pulp. That trait is also inherited in the hybrids with *Soubre* as a female parent.

**Table 3:**
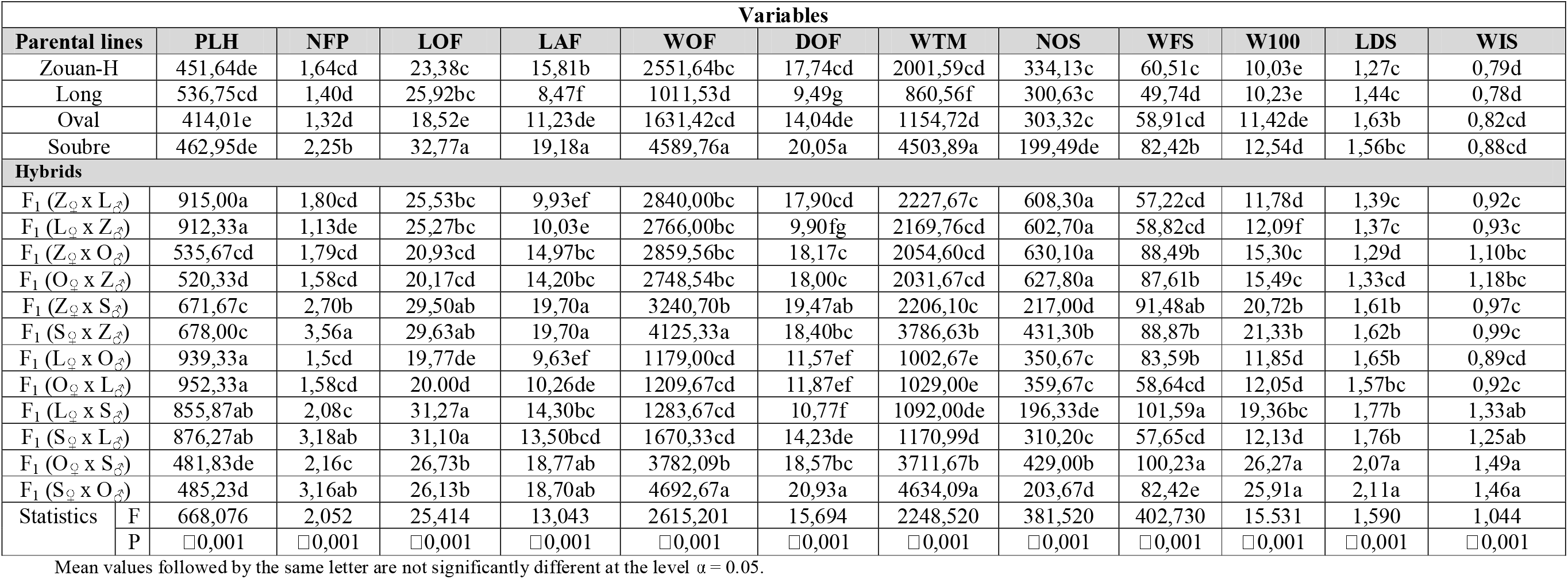
Means of plant height, fruit- and seed-yield traits of the parental lines and the hybrid combinations of *Cucurbita moschata D* used in this study.

**Figure 1.**
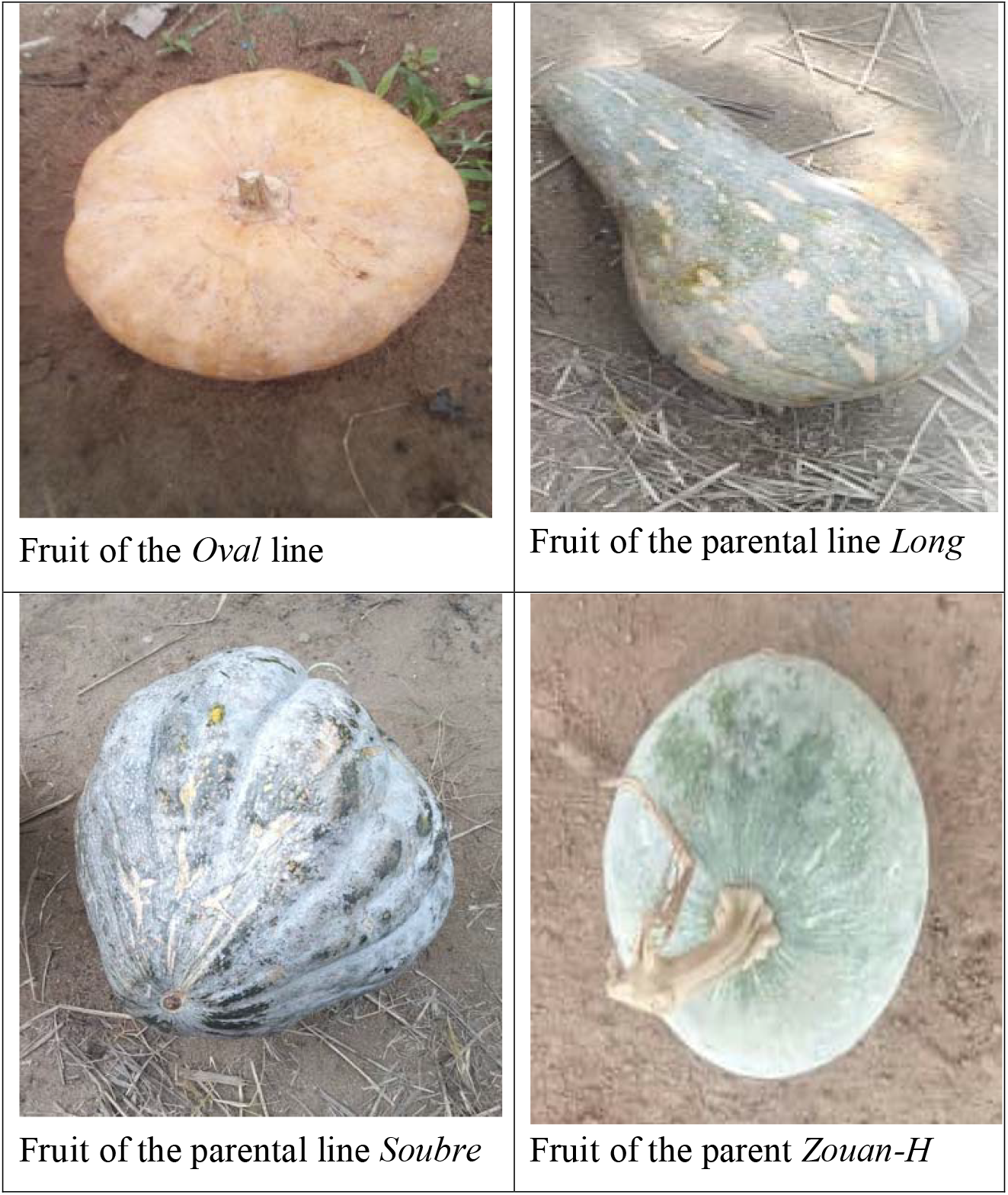
Diversity of the fruits of the parental lines of *Curcubita moschata* D.

The parental lines *Zouan-H, Long* and *Oval* have significantly higher number of seeds per fruit. But the seeds are smaller and lighter. The hybrids produced from these three parents also have the highest number of seeds. In particular, *Z*_♀_ x *L*_♂_, *L*_♀_ x *Z*_♂,_ *Z*_♀_ x *O*_♂_ and *O*_♀_ x *Z*_♂_ have doubled the number of seeds per fruit compared to their parents. Hybrids produced from those three parents have smaller seeds that are lighter. The *Soubre* parent has a smaller number of seeds that are comparatively bigger and heavier. In most cases, hybrids involving *Soubre* as a parent have smaller number of seeds per fruit, except *O*_♀_ x *S*_♂_ and *S*_♀_ x *Z*_♂_. But, they have heavier and bigger seeds. In most cases, the hybrids outperformed the parents. The hybrids *S*_♀_ x *O*_♂_ and *O*_♀_ x *S*_♂_ more than doubled their 100-seed weight compared to either parent.

### 3.2. Heterosis and Values of the Potency Ratio for the Different Traits

#### 3.2.1. Estmates of heterosis in the F1 Hybrids

The estimates of heterosis expressed as a percent increase (or decrease) with respect to the average of the two parents (*mph*) and the better parent (*bph*) are presented in Tables 4 and 5, respectively.

**Table 4:**
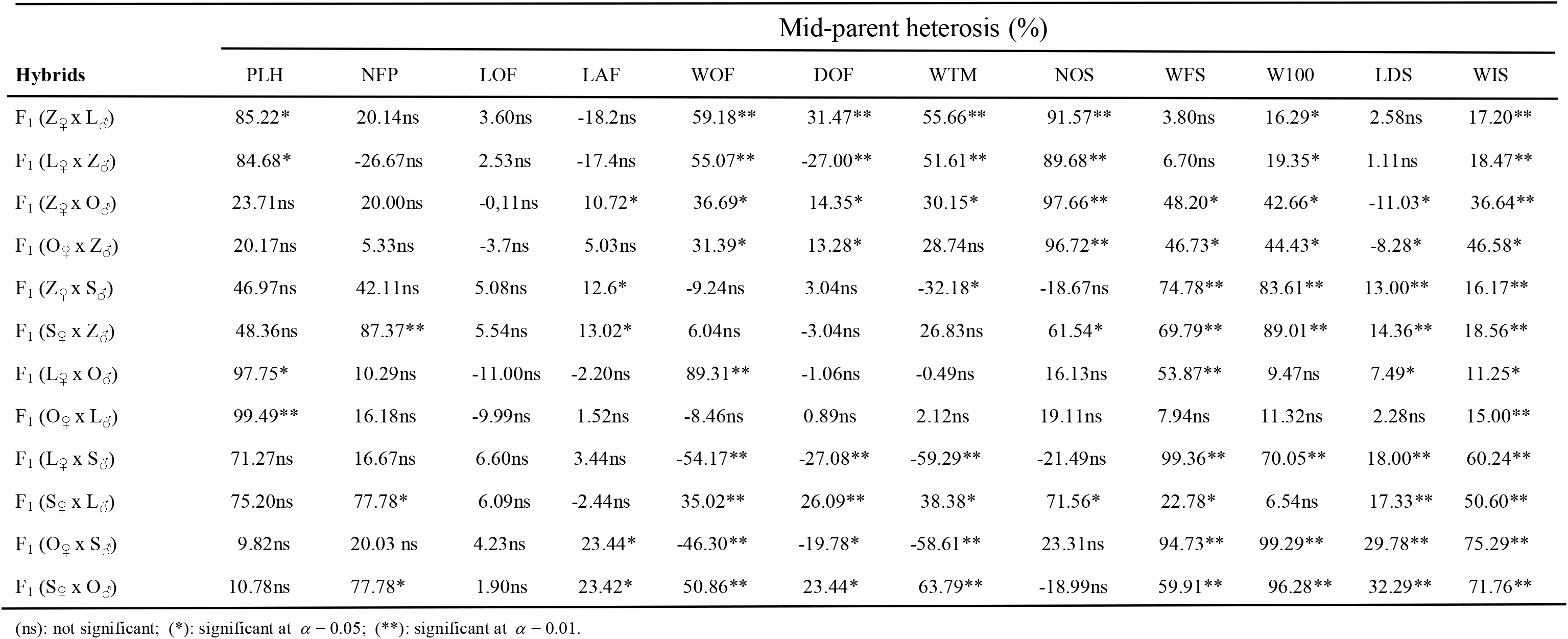
Mid-parent heterosis (in percentages) observed on fruit- and seed-yield traits and plant height of the hybrids.

| Hybrids | Mid-parent heterosis (%) |  |  |  |  |  |  |  |  |  |  |  |
| --- | --- | --- | --- | --- | --- | --- | --- | --- | --- | --- | --- | --- |
|  | PLH | NFP | LOF | LAF | WOF | DOF | WTM | NOS | WFS | W100 | LDS | WIS |
| F<sub>1</sub> (Z<sub>♀</sub> x L<sub>♂</sub>) | 85.22* | 20.14ns | 3.60ns | -18.2ns | 59.18** | 31.47** | 55.66** | 91.57** | 3.80ns | 16.29* | 2.58ns | 17.20** |
| F<sub>1</sub> (L<sub>♀</sub> x Z<sub>♂</sub>) | 84.68* | -26.67ns | 2.53ns | -17.4ns | 55.07** | -27.00** | 51.61** | 89.68** | 6.70ns | 19.35* | 1.11ns | 18.47** |
| F<sub>1</sub> (Z<sub>♀</sub> x O<sub>♂</sub>) | 23.71ns | 20.00ns | -0.11ns | 10.72* | 36.69* | 14.35* | 30.15* | 97.66** | 48.20* | 42.66* | -11.03* | 36.64** |
| F<sub>1</sub> (O<sub>♀</sub> x Z<sub>♂</sub>) | 20.17ns | 5.33ns | -3.7ns | 5.03ns | 31.39* | 13.28* | 28.74ns | 96.72** | 46.73* | 44.43* | -8.28* | 46.58* |
| F<sub>1</sub> (Z<sub>♀</sub> x S<sub>♂</sub>) | 46.97ns | 42.11ns | 5.08ns | 12.6* | -9.24ns | 3.04ns | -32.18* | -18.67ns | 74.78** | 83.61** | 13.00** | 16.17** |
| F<sub>1</sub> (S<sub>♀</sub> x Z<sub>♂</sub>) | 48.36ns | 87.37** | 5.54ns | 13.02* | 6.04ns | -3.04ns | 26.83ns | 61.54* | 69.79** | 89.01** | 14.36** | 18.56** |
| F<sub>1</sub> (L<sub>♀</sub> x O<sub>♂</sub>) | 97.75* | 10.29ns | -11.00ns | -2.20ns | 89.31** | -1.06ns | -0.49ns | 16.13ns | 53.87** | 9.47ns | 7.49* | 11.25* |
| F<sub>1</sub> (O<sub>♀</sub> x L<sub>♂</sub>) | 99.49** | 16.18ns | -9.99ns | 1.52ns | -8.46ns | 0.89ns | 2.12ns | 19.11ns | 7.94ns | 11.32ns | 2.28ns | 15.00** |
| F<sub>1</sub> (L<sub>♀</sub> x S<sub>♂</sub>) | 71.27ns | 16.67ns | 6.60ns | 3.44ns | -54.17** | -27.08** | -59.29** | -21.49ns | 99.36** | 70.05** | 18.00** | 60.24** |
| F<sub>1</sub> (S<sub>♀</sub> x L<sub>♂</sub>) | 75.20ns | 77.78* | 6.09ns | -2.44ns | 35.02** | 26.09** | 38.38* | 71.56* | 22.78* | 6.54ns | 17.33** | 50.60** |
| F<sub>1</sub> (O<sub>♀</sub> x S<sub>♂</sub>) | 9.82ns | 20.03 ns | 4.23ns | 23.44* | -46.30** | -19.78* | -58.61** | 23.31ns | 94.73** | 99.29** | 29.78** | 75.29** |
| F<sub>1</sub> (S<sub>♀</sub> x O<sub>♂</sub>) | 10.78ns | 77.78* | 1.90ns | 23.42* | 50.86** | 23.44* | 63.79** | -18.99ns | 59.91** | 96.28** | 32.29** | 71.76** |

(ns): not significant; (\*): significant at $\alpha = 0.05$ ; (\*\*): significant at $\alpha = 0.01$ .

**Table 5:** Best-parent heterosis observed with the agronomic, fruit- and seed-yield traits of the hybrids.

| Hybrids | Best-parent heterosis (%) |  |  |  |  |  |  |  |  |  |  |  |
| --- | --- | --- | --- | --- | --- | --- | --- | --- | --- | --- | --- | --- |
|  | PLH | NFP | LOF | LAF | WOF | DOF | WTM | NOS | WFS | W100 | LDS | WIS |
| F<sub>1</sub> (Z<sub>♀</sub> x L<sub>♂</sub>) | 70.47ns | 9.76ns | -1.50ns | -37.20* | 11.01ns | 0.94ns | 11.30ns | 81.97** | -5.44ns | 15.15* | -3.47ns | 16.46** |
| F<sub>1</sub> (L<sub>♀</sub> x Z<sub>♂</sub>) | 69.97ns | -32.93ns | -2.50ns | -36.56* | 8.06ns | -44.05** | 8.40ns | 80.17** | -2.79ns | 18.18* | -4.86ns | 17.72** |
| F<sub>1</sub> (Z<sub>♀</sub> x O<sub>♂</sub>) | 18.83ns | 9.76ns | -10.5ns | -5.30ns | 12.20ns | 3.25ns | 2.62ns | 88.55** | 46.24* | 33.98* | -20.86** | 34.15** |
| F<sub>1</sub> (O<sub>♀</sub> x Z<sub>♂</sub>) | 15.21ns | -3.66ns | -13.73ns | -10.20ns | 7.69ns | 1.47ns | 1.50ns | 87.65** | 44.79* | 35.64* | -18.40** | 43.90** |
| F<sub>1</sub> (Z<sub>♀</sub> x S<sub>♂</sub>) | 45.08ns | 20.00ns | -10.00ns | 2.71ns | -29.39* | -2.89ns | -51.02** | -35.06ns | 51.18** | 65.23** | 3.02ns | 10.23* |
| F<sub>1</sub> (S<sub>♀</sub> x Z<sub>♂</sub>) | 46.45ns | 58.22ns | -9.58ns | 2.71ns | -17.01* | -8.23ns | -8.41ns | 28.99ns | 46.87* | 70.09** | 3.40ns | 12.50* |
| F<sub>1</sub> (L<sub>♀</sub> x O<sub>♂</sub>) | 75.32ns | 7.14ns | -23.73ns | -14.20ns | -27.73* | -18.13* | -13.17ns | 15.61ns | 41.89* | 3.77ns | 1.23ns | 8.54* |
| F<sub>1</sub> (O<sub>♀</sub> x L<sub>♂</sub>) | 77.43ns | 12.86ns | -22.80ns | -10.95ns | -26.09* | -15.27ns | -10.89ns | 18.58ns | -0.46ns | 5.52ns | -3.68ns | 12.19* |
| F<sub>1</sub> (L<sub>♀</sub> x S<sub>♂</sub>) | 59.29ns | -7.56ns | -4.58ns | -25.40* | -72.10 ** | -46.28** | -75.75** | -34.39ns | 89.24** | 54.34** | 13.46** | 51.14** |
| F<sub>1</sub> (S<sub>♀</sub> x L<sub>♂</sub>) | 63.20ns | 41.33ns | -5.10ns | -29.61* | -17.59* | -7.28ns | -20.48ns | 42.70* | 15.90* | -3.27ns | 12.82* | 42.05** |
| F<sub>1</sub> (O<sub>♀</sub> x S<sub>♂</sub>) | 3.90ns | -4.18 ns | -18.40ns | -2.10ns | -64.11 ** | -29.03** | -74.00** | 2.20ns | 70.14** | 89.49** | 26.99** | 69.32** |
| F<sub>1</sub> (S<sub>♀</sub> x O<sub>♂</sub>) | 4.81ns | 42.22ns | -20.3ns | -2.55ns | 2.24ns | 4.25ns | 2.89ns | -32.85ns | 39.91* | 86.62** | 29.45** | 65.91** |

(ns): not significant; (\*): significant at $\alpha = 0.05$ ; (\*\*): significant at $\alpha = 0.01$ .

Mid-parent heterosis was found for all measured traits in the crosses, except the length of the fruit. For the character plant height, significant mid-parent heterosis was observed with hybrids involving the parents *Long* and *Zouan-H*, and the parents *Long* and *Ova*l. Regarding the number of fruits per plant, mid-parent heterosis was significant only in hybrids where *Soubre* is the female parent. None of the estimated better-parent heterosis was significant for the number of fruits per plant. For the length of the fruit, none of the estimated heterosis was significant, whether computed on the basis of the average of the two parents or the better parent. Significant mid-parent heterosis was observed in the hybrids from the crosses between the parental lines *Soubre* and *Ovale, Soubre* and *Zouan-H*, and in the hybrid *Z*_♀_ x *O*_♂_ for the character width of the fruit. For that character, the significant estimates of heterosis based on the best parent were negative. They were observed in the hybrids involving the parents *Zouan-H* and *Long*, and *Soubre* and *Long*, whether used as male or female parent. For the diameter of the fruit, significant estiamtes of mid-parent heterosis were observed in the crosses *Z*_♀_ x *L*_♂_, *S*_♀_ x *L*_♂_, and *S*_♀_ x *O*_♂_. The reciprocal crosses yielded significant negative estimates of mid-parent heterosis. Also, significant estimates of heterosis were observed with the hybrids from the cross *Z*_♀_ x *O*_♂_ and its reciprocal. All significant estimates of heterosis computed on the basis of the better parent were negative. They were observed in the hybrids *L*_♀_ x *Z*_♂_, *L*_♀_ x *O*_♂_, *L*_♀_ x *S*_♂_, and *O*_♀_ x *S*_♂_. For the weight of the pulp, all estimates of mid-parent heterosis were significant except in the following hybrids *O*_♀_ x *Z*_♂_, *S*_♀_ x *Z*_♂_, *L*_♀_ x *O*_♂_, and *O*_♀_ x *L*_♂_. However, the heterosis estimated on the basis of the better parent was not significant for the same trait in all hybrids except *Z*_♀_ x *S*_♂_, *L*_♀_ x *S*_♂_, and *O*_♀_ x *S*_♂_. For the character number of seeds per fruit, significant estimates of heterosis computed with either method were seen in the crosses involving *Zouan-H* and *Long* and *Zouan-H* and *Oval* and their reciprocals. Mid-parent heterosis was also significant in the hybrids *S*_♀_ x *Z*_♂_ and *S*_♀_ x *L*_♂_ and better-parent heterosis was significant in the latter hybrid. With the weight of fresh seeds per fruit, heterosis whether estimated with the average of the two parents or with the better parent, was significant in all crosses except *Z*_♀_ x *L*_♂_, *L*_♀_ x *Z*_♂_ and *O*_♀_ x *L*_♂_. And significant heterosis was found for the 100-seed weight in all hybrids except *L*_♀_ x *O*_♂_, *O*_♀_ x *L*_♂_ and *S*_♀_ x *L*_♂_. That observation applied to the two methods of estimation. Regarding the character length of the dry seed, mid-parent heterosis was significant in all hybrids except those from crosses involving *Zouan-H* and *Long*, and in the hybrid *O*_♀_ x *L*_♂._ For the same character, estimation of heterosis based on the better parent was significant only in crosses where the parents were *Zouan-H* and *Oval, Long* and *Soubre*, and *Oval* and *Soubre*. In hybrids from the crosses involving *Zouan-H* and *Oval*, better-parent heterosis was negative. For the width of the dry seed, heterosis computed with either method was significant in all hybrids.

These results implied that the different hybrids exhibited varied performances according to the traits observed.

#### 3.2.2. Effect of Gene Dominance

The effects of gene dominance are determined by the values of the potency ratio. They are reported in Table 6. In the absence of a significance test, values were interpreted as they were. We did not assume a value -0.99 to equate -1.00 or a value 0.05 to equate 0.00, without statistical support. Doing so would give totally different interpretations of gene expression in many cases of the quantitative analysis.

**Table 6:** Relative potency ratio for the fruit- and seed-yield traits and plant height of the hybrids.

| Hybrids | Relative potency ratio |  |  |  |  |  |  |  |  |  |  |  |
| --- | --- | --- | --- | --- | --- | --- | --- | --- | --- | --- | --- | --- |
|  | PLH | NFP | LOF | LAF | WOF | DOF | WTM | NOS | WFS | W100 | LDS | WIS |
| F<sub>1</sub> (Z<sub>♀</sub> x L<sub>♂</sub>) | 9.89 | 2.50 | 0.69 | -0.60 | 1.37 | 1.04 | 1.40 | 17.35 | 0.39 | 16.50 | 2.43 | 27.00 |
| F<sub>1</sub> (L<sub>♀</sub> x Z<sub>♂</sub>) | 9.83 | -0.33 | 0.49 | -0.57 | 1.28 | -0.90 | 2.59 | 16.99 | 0.69 | 9.80 | 0.18 | 29.00 |
| F<sub>1</sub> (Z<sub>♀</sub> x O<sub>♂</sub>) | 5.47 | 1.81 | -0.10 | 0.63 | 1.67 | 1.23 | 1.12 | 10.10 | 35.98 | 6.58 | 0.89 | 19.67 |
| F<sub>1</sub> (O<sub>♀</sub> x Z<sub>♂</sub>) | 4.64 | 0.08 | -0.32 | 0.30 | 1.43 | 1.14 | 1.07 | 20.01 | 34.88 | 6.86 | -0.30 | 25.00 |
| F<sub>1</sub> (Z<sub>♀</sub> x S<sub>♂</sub>) | 37.96 | 2.29 | 0.30 | 1.30 | -0.32 | 0.50 | -0.84 | -0.74 | 4.79 | 7.52 | 1.10 | 3.00 |
| F<sub>1</sub> (S<sub>♀</sub> x Z<sub>♂</sub>) | 39.61 | 5.30 | 0.33 | 1.31 | 0.21 | -0.43 | 0.70 | 2.44 | 4.47 | 8.00 | 1.58 | 3.44 |
| F<sub>1</sub> (L<sub>♀</sub> x O<sub>♂</sub>) | 7.56 | 3.51 | -0.66 | -0.20 | 0.46 | 0.09 | -0.03 | 36.20 | 6.38 | 1.72 | 1.21 | 4.50 |
| F<sub>1</sub> (O<sub>♀</sub> x L<sub>♂</sub>) | 7.78 | 3.26 | -0.60 | 0.10 | -0.36 | 0.05 | 0.15 | 42.90 | 0.94 | 2.06 | 0.37 | 6.00 |
| F<sub>1</sub> (L<sub>♀</sub> x S<sub>♂</sub>) | 8.90 | 0.66 | 0.56 | 0.10 | -0.85 | -0.76 | -0.87 | -0.98 | 51.85 | 6.90 | 4.50 | 10.00 |
| F<sub>1</sub> (S<sub>♀</sub> x L<sub>♂</sub>) | 10.19 | 3.19 | 0.51 | -0.11 | 0.55 | 0.72 | 0.57 | 3.54 | 3.84 | 0.65 | 4.33 | 8.40 |
| F<sub>1</sub> (O<sub>♀</sub> x S<sub>♂</sub>) | 1.76 | 0.77 | 0.15 | 0.90 | -0.97 | -0.94 | -0.99 | 1.13 | 6.61 | 25.52 | 13.57 | 2.00 |
| F<sub>1</sub> (S<sub>♀</sub> x O<sub>♂</sub>) | 1.93 | 2.96 | 0.07 | 0.88 | 1.07 | 1.29 | 1.08 | -0.92 | 4.19 | 24.88 | 14.71 | 20.33 |

The gene(s) controlling plant height exhibited a super-dominance in all the hybrids examined because the potency ratio was greater than one (*P* > 1) in all the crosses of the inbred lines of *Cucurbita moschata* used in this study. For the number of fruits per plant, a partial dominance was observed in the following F1 hybrids *L*_♀_ x *Z*_♂_, *O*_♀_ x *Z*_♂_, *L*_♀_ x *S*_♂_, and *O*_♀_ x *S*_♂_ because -1 < *P* < 1 and *P* ≠ 0, and a super-dominance in the expression of the character in all the other hybrids. The negative value of the potency ratio observed in the F1 hybrid *L*_♀_ x *Z*_♂_ indicated a reduction in the number of fruits per plant for that hybrid. Partial dominance was observed in the expression of the length of the fruits in all hybrids. For F_1_ (*Z*_♀_ x *O*_♂_) and F_1_ (*L*_♀_ x *O*_♂_) and their respective reciprocals, the values of the potency ratio were negative, indicating a relative reduction of the length of the fruits in those hybrids. The F1 hybrids issued from the cross *Z*_♀_ x *S*_♂_ and its reciprocal showed a super-dominance of the gene(s) determining the width of the fruits in those hybrids. In all the other hybrid combinations, partial dominance was observed in the expression of the width of the fruit, and the values of the relative potency ratio were negative for the following F1 hybrids *Z*_♀_ x *L*_♂_, *L*_♀_ x *Z*_♂_, *L*_♀_ x *O*_♂_, and *S*_♀_ x *L*_♂_. For the weight of the fruit, a super-dominance in the expression of the trait was observed in the F1 hybrids *Z*_♀_ x *L*_♂_, *L*_♀_ x *Z*_♂_, *Z*_♀_ x *O*_♂_, *O*_♀_ x *Z*_♂_, and *S*_♀_ x *O*_♂_. All the other hybrids exhibited partial dominance, and the values of the potency ratio were negative for the following F1 hybrids *Z*_♀_ x *S*_♂_, *O*_♀_ x *L*_♂_, *L*_♀_ x *S*_♂_, and *O*_♀_ x *S*_♂_. Expression of the diameter of the fruit varied too, according to the hybrids. A super-dominance in the expression of the diameter of the fruit was observed in the F_1_ (*Z*_♀_ x *L*_♂_), F_1_ (*Z*_♀_ x *O*_♂_), F_1_ (*O*_♀_ x *Z*_♂_), and F_1_ (*S*_♀_ x *O*_♂_), and partial dominance for the character in all the other F1 hybrids. Four F1 hybrids had a negative value of the relative potency ratio for the diameter of the fruit. For the weight of the pulp, the expression of the trait varied according to the hybrids, but in a different pattern. In the F1 hybrids from the crosses involving *Zouan-H* and *Long, Zouan-H* and *Oval*, their respective reciprocals and the F_1_ (*S*_♀_ x *O*_♂_), the genes that determined the weight of the pulp were super-dominant. In all the other hybrids, partial dominance was observed in the expression of the character with some cases of negative value of the potency ratio. The expressions of the examined seed-related traits were mostly super-dominant. For the number of seeds per fruit, super-dominance was observed in the determination of the character in all F1 hybrids except F_1_ (*Z*_♀_ x *S*_♂_), F_1_ (*L*_♀_ x *S*_♂_), and F_1_ (*S*_♀_ x *O*_♂_) where -1 < *P* < 0. The genes governing the weight of fresh seeds were super-dominant in all F1 hybrids except F_1_ (*Z*_♀_ x *L*_♂_), F_1_ (*L*_♀_ x *Z*_♂_), and F_1_ (*O*_♀_ x *L*_♂_) where partial dominance was observed. A super-dominance was observed in all hybrids for the 100-seed weight except F_1_ (S_♀_ x L_♂_) where we observed partial dominance. For the length of the dry seed, partial dominance was observed in the F_1_ (*L*_♀_ x *Z*_♂_), F_1_ (*Z*_♀_ x *O*_♂_), F_1_ (*O*_♀_ x *Z*_♂_), and F_1_ (*O*_♀_ x *L*_♂_). In all the other hybrid combinations, the expression of the length of the dry seed was characterized by a super-dominance of the long seed. And the width of the seed was characterized by a super-dominance of the wide seed in all hybrids studied.

#### 3.2.3 Analysis of variance of the *gca*’s and *sca*’s of the traits

The examined traits significantly differed according to blocks (Table 7). Across all the blocks, we observed highly significant *gca* effects of the female parents for all traits studied, indicating a very large variation of the *gca* effects of the inbred lines when used as females. When used as male parents, the *gca* effects were significant for all the traits except plant height, length of the fruit, width of the fruit and length of the seed. For all the traits, the variation in *gca* of the female parent was far greater than that of the male parent. Highly significant *sca* effects were observed for all the traits studied. The reciprocal effects were highly significant for weight and diameter of the fruit, weight of the pulp, number of seeds per fruit, weight of the fresh seeds and 100-seed weight, indicating strong maternal effects on those traits. The general predictive ratio is given as 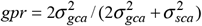 . A value of *gpr* = 2/3 indicates equal contribution of additive and non-additive gene effects in the determination of a trait. A value of *gpr* < 2/3 means the predominance of non-additive gene effects over the additive ones in the expression of the trait. And a value of *gpr* > 2/3 means that additive gene effects predominantly determine the trait. Especially, a *gpr* closer to 1 means only additive gene effects control the expression of the trait. In this study, the general predictive ratio was greater than two-third for all the traits except weight of the fresh seeds and width of the dry seed. We may infer from those observations that most of the character under this study are predominantly determined by the effects of additive gene. But, weight of the fresh seeds and width of the dry seed are mostly controlled by non-additive gene effects.

**Table 7:** Analysis of variance of the gca’s and sca’s effects based on the linear diallel model, and general predictive ratio of the fruit- and seed-yield traits and plant height of four inbred lines and their hybrid combinations of *Cucurbita moschata*.

| SOV | df | Mean Square |  |  |  |  |  |  |  |  |  |  |  |
| --- | --- | --- | --- | --- | --- | --- | --- | --- | --- | --- | --- | --- | --- |
|  |  | PLH | NFP | LOF | LAF | WOF | DOF | WTM | NOS | WFS | W100 | LDS | WIS |
| Block | 2 | 3961432** | 6.198** | 2854.2** | 933.7** | 261077454** | 2597.1** | 223039259** | 363343** | 24797** | 243.2** | 0.133* | 0.076ns |
| GCA(female) | 3 | 8706682** | 153.516** | 10970.5** | 8056.6** | 431617966** | 4777.5** | 468478919** | 5827578** | 29220** | 6228.8** | 20.709** | 7.501** |
| GCA(male) | 3 | 11451ns | 60.834** | 5.8ns | 7.5ns | 239998329** | 2560.2** | 397639672** | 1045979** | 41302** | 469.6** | 0.008ns | 0.178** |
| SCA | 6 | 7014551** | 38.308** | 248.3** | 389.8** | 34229753** | 109.7** | 19207823** | 2821242** | 57108** | 4376.9** | 6.326** | 10.520** |
| Reciprocal | 3 | 5135ns | 1.527ns | 13.1ns | 16.0ns | 42467338** | 517.0** | 17294741** | 962334** | 10541** | 482.9** | 0.045ns | 0.066ns |
| Residuals | 1710 | 98411 | 1.389 | 57.5 | 11.8 | 2131174 | 18.7 | 2110005 | 27171 | 664 | 18.0 | 0.039 | 0.031 |
| *gpr* |  | 0.713 | 0.889 | 0.989 | 0.976 | 0.962 | 0.989 | 0.979 | 0.918 | 0.506 | 0.740 | 0.867 | 0.588 |

**Table 8:** Estimation of the general combining abilities (gca) of the parental lines and the specific combining abilities (sca) of the F1 hybrids for the fruit- and seed-yield traits and plant height of *Cucurbita moschata* materials used in this study.

|  | PLH | NFP | LOF | LAF | WOF | DOF | WTM | NOS | WFS | W100 | LDS | WIS |
| --- | --- | --- | --- | --- | --- | --- | --- | --- | --- | --- | --- | --- |
| Parental lines | gca of parents |  |  |  |  |  |  |  |  |  |  |  |
| Zouan-H | -25.93** | -0.07* | -0.69** | 0.76** | 302.82** | 1.44** | 99.21ns | 91.54** | 1.07ns | -0.88** | -0.19* | -0.19* |
| Long | 147.48** | -0.29** | 0.18ns | -3.72** | -729.70** | -3.17** | -633.79** | 21.99** | -8.49* | -3.08** | -0.04ns | -0.04ns |
| Oval | -75.21** | -0.25** | -4.06** | -0.67** | -412.43** | -0.36* | -464.96** | 16.58** | 4.23* | -0.66** | 0.06ns | 0.06ns |
| Soubre | -46.34** | 0.62** | 4.57** | 3.62** | 839.31** | 2.09** | 999.54** | -108.12** | 3.20* | 3.30** | 0.16* | 0.17* |
| (SE) | (39) | (0.05) | (0.37) | (0.18) | (124.15) | (0.36) | (235.99) | (6.94) | (2.39) | (0.44) | (0.01) | (0.009) |
| Mating | sca of crosses |  |  |  |  |  |  |  |  |  |  |  |
| Z ⊗ L | 124.16** | -0.22* | 1.49** | -1.32** | 614.75** | -0.09ns | 480.23** | 120.09** | -7.67* | 0.36* | 0.03* | 0.03* |
| Z ⊗ O | -38.83* | -0.04ns | -0.10ns | 0.23ns | 297.98** | 1.28** | 155.57ns | 151.01** | 9.63** | 0.06ns | -0.15* | -0.15* |
| Z ⊗ S | 79.14** | 0.54** | 0.27ns | 1.06** | 243.60** | -0.32ns | -186.10ns | -40.79* | 12.79** | 3.06** | -0.05* | -0.05* |
| L ⊗ O | 205.60** | 0.34** | -1.64** | -0.06ns | -278.65** | -1.47** | -138.43ns | -42.78* | 2.26ns | -1.18* | -0.04* | -0.04* |
| L ⊗ S | 96.40** | 0.26** | 1.01** | -0.20ns | 391.90** | 0.02ns | -276.93** | 27.42* | 11.79** | -0.02ns | -0.05* | -0.05* |
| O ⊗ S | -63.24** | 0.24** | 0.51 | 1.52** | 339.50** | 0.13ns | 151.24ns | -21.00* | 10.77** | 6.57** | 0.27* | 0.27* |
| (SE) | (35.85) | (0.11) | (0.69) | (0.33) | (226.67) | (0.67) | (235.99) | (12.67) | (4.38) | (0.82) | (0.02) | (0.02) |

(ns) means not significant; (\*) means significant at $\alpha = 0.05$ ; (\*\*) means significant at $\alpha = 0.01$ ; (⊗) stands for direct cross and reciprocal; (SE) means standard error of the gca or sca effects.

In particular, traits related to the fruit have a general predictive ratio greater than 0.9 and very close to 1, meaning that they are almost solely determined by additive gene effects.

The estimates of general combining abilities varied across parental lines as well as the traits, with negative and positive values (Table 8). In general, negative *gca* effects would be desirable for plant height. We would want the plant to complete its vegetative growth early enough in order to allocate larger proportion of dry matter to the development of the fruits and the seeds. Positive *gca* effects would be preferred for fruit- and seed-related traits because they are the commonly harvestable organs that determine yield. The parental line *Soubre* had negative *gca* effects for plant height and positive *gca* effects for all the fruit- and seed-related traits except the number of seeds per fruit. The parent *Soubre* would be suitable for developing a shorter *C. moschata* hybrid with higher number of large fruits, and large seeds. The parent *Zouan-H* has negative *gca* effects for plant height, and positive *gca* effects for fruit- and seed-related traits except the number of fruits per plant and fruit length. Both *Soubre* and *Zouan-H* have very high estimates of *gca* effects on the weight of the fruit and pulp mass. If interest lies in a longer vegetative growth with reduced fruit- and seed-yield, the inbred line *Long* would be preferred as it has very high *gca* effects for plant height and negative or non-significant *gca* effects for fruit- and seed-related traits, except number of seeds per fruit. The parental line *Oval* would be suitable for reducing plant height and increasing the number of seeds per fruit. It has significant negative *gca* effects for plant height, and positive *gca* effects for number of seeds per fruit.

Based on the *sca* effects, we may affirm that crosses involving the parental line *Long* would result in hybrids with increased plant height. Crosses between *Zouan-H* and *Long* would increase fruit weight, pulp weight, and number of seeds per fruit, along with an increased plant height. And crosses between *Zouan-H* and *Oval* would produce hybrids with significantly reduced plant height, increased fruit weight and increased number of seeds per fruit. Combination of crosses involving the parent *Soubre* would result in hybrids with increased fruit yield. And hybrids from crosses involving *Oval* and *Soubre* would have reduced plant height, increased fruit weight and increased seed weight.

Random samples from each inbred line in each of the three blocks were used to develop a classification model, using the *gca* of the parental lines. Based on the within-group sum-of-squares, a three-group model was found to minimize the error variance. Figure 2 is the unrooted classification tree that assembled the parental lines in three groups based on their *gca* effects, each group with a distinctive color. The first group is identified with the green color and it assembled the parental lines *Soubre*. The *Soubre* line used as a parent had a very high general combining ability effect on fruit traits such as weight of the fruit, diameter of the fruit, length of the fruit, weight of the pulp, and number of fruits per plant. It also has a high positive *gca* effect on the one-hundred seed weight. The second group is identified with the orange color and includes the parental lines *Zouan-H*. It is characterized by a significantly high gca effect on the weight of the fruit, the diameter of the fruit, and the number of seeds per fruit but the seeds have a reduced size. The third group, identified with the red color, includes the parental lines *Long* and *Oval*. They both have significant gca effects on the number of seeds per fruit, and reduced seed size. The parental line *Long* is mostly characterized by an extended vegetative growth.

**Figure 2:**
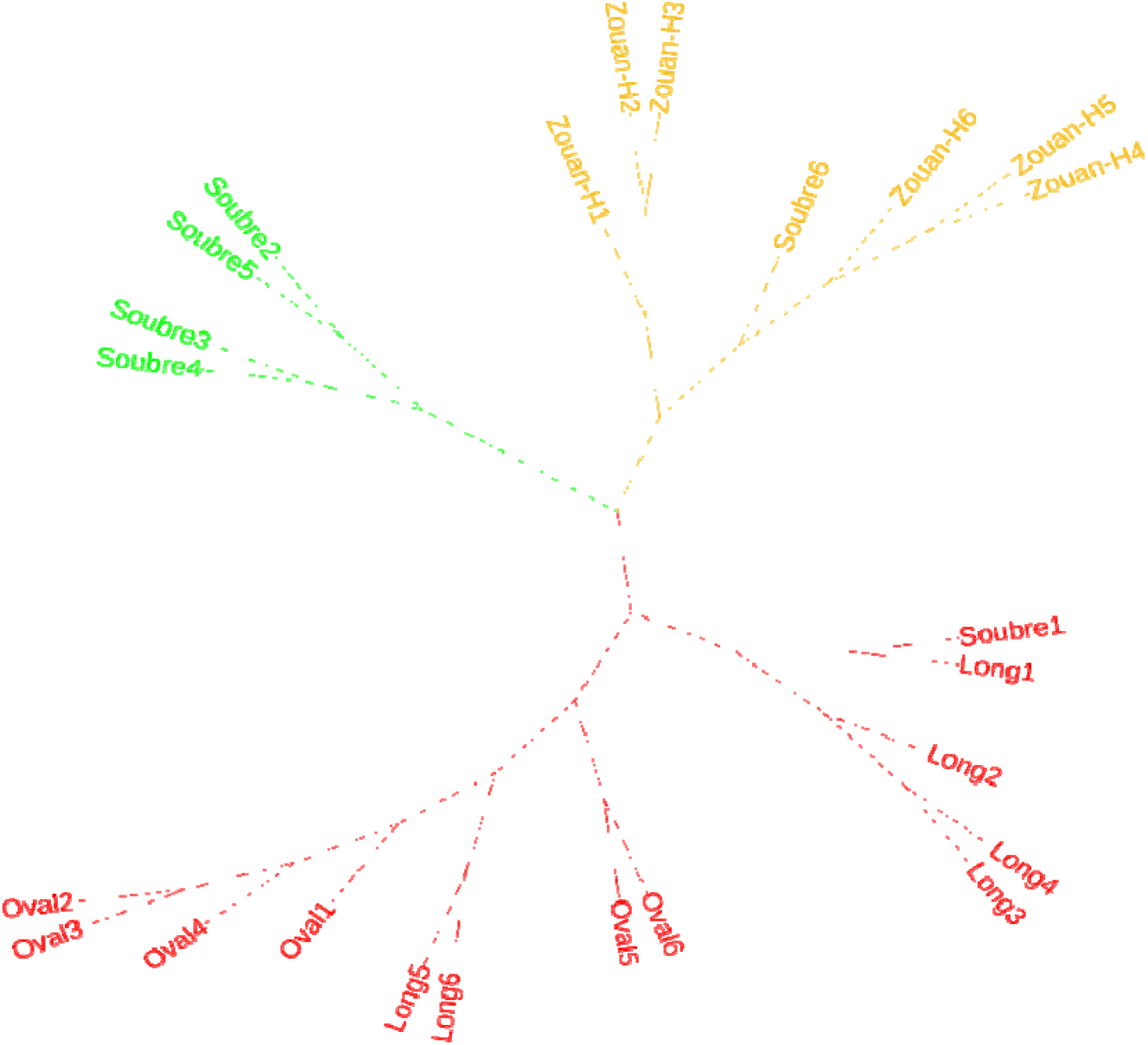
Unrooteed classification tree of the parental lines, based on their general combining ability effects.

Hence, *Soubre* would be a promising line in a breeding program where improvement of fruit yield is the objective. The parent *Long* would be promising in a program to develop hybrids for extended vegetative development. For improvement of the seed traits, the parental line *Zouan-H* or *Oval* could serve as a female parent if an increase in the number of seeds per fruit is the objective. The parent *Soubre* could also serve as a female parent if the objective is to increase the size of the seeds.

## 4. Discussion

The evaluation of the parental lines with respect to the fruits, the seed traits and plant height showed that the lines *Soubre, Oval, Long*, and *Zouan-H* have very contrasting characters, and pointed out the existence of a large genetic diversity in the butternut squash germplasm. In fact, the genetic diversity of this species is considerable in terms of the shapes, the forms and the sizes of the fruits and the seeds, their growth cycle, and their degree of resistance to viral diseases and conservation [24]. We should mention that the phenotypic variability in these parental lines has already been described [25]. The phenotypic variability observed in the four lines can be used in selection program for the genetic improvement of the traits of interest in *Cucurbita moschata* D. [26, 27].

The study of the performances of the parents and their F1 hybrid combinations revealed significant differences between the genotypes for each of the character evaluated in this work. Some F1 hybrids obtained much better performances than others for each trait studied. The hybrids could be proposed as candidates in breeding programs aimed at improving traits where theses hybrids have recorded their best performances [28].

Significant positive heterosis effects were observed in different hybrids for several traits. This hybrid performance could be a consequence of the large genetic distance separating the different parental lines from each other [29]. Hybrids resulting from genetically distant parents express a heterosis effect [30]. And hybrid vigor is based on the complementation of the gametic contributions of the parents by favorable dominant genes [31] and therefore, the expression of the heterosis effect could be linked in part to the non-additive action of genes, dominance and epistasis. The detection of heterosis in a species is an important step for the improvement of the species. Indeed, the expression of heterosis by hybrids for a given trait indicates their potential to produce superior cultivars through the selection of transgressive segregants in segregating populations [32, 33]. Heterosis breeding is a potential tool to improve the quantity, quality and productivity of bitter gourd [34], which cannot be done by traditional methods. Analysis of variance of the *gca* and *sca* revealed significant general and specific combining ability effects for the evaluated traits. These results suggested the involvement of additive and non-additive gene actions in the expression of the characters. Estimates of the general predictive ratio, 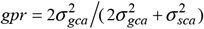, for each trait varied between 0.506 and 0.989 with most estimates being greater than 0.900. A *gpr* estimate of 2/3 is the indication of equal variance of the *gca* and the *sca* in the distribution of the variances among the components. For the traits studied in this experiment, the *gpr* estimates were greater than 2/3 for all traits except the weight of the fresh seeds and the width of the dry seed. This indicated the preponderance of additive gene effects in the genetic control of most of the traits studied. The average performance of a genotype in a series of hybrid combinations will be most relevant in the design of a breeding procedure in cultivar development. If a particular cultivar has a high *gca* for a trait, it means that the cultivar would be a valuable parent in the breeding program to improve that trait [35]. In our study, *Soubre* is the best general combiner for a large number of characters studied with the exception of the number of seeds per fruit and plant height. The lines *Oval, Long* and *Zouan-H* stand out as the best overall combiners for number of seeds per fruit. *Long* is the best general combiner for plant height. Thus, each of the four parental lines could be used in a selection program to improve the traits of interest in *Cucurbita moschata* D. Also, the significant *gca* values for a given trait revealed that the selection and hybridization methods would lead to interesting genetic improvement for the trait, due to the accumulation of desirable and favorable alleles from both parents in the targeted genotype [28].

The best combination in terms of *gca* effects consistently involved one or two best overall combiners as parents. In other words, most crosses resulting in high *gca* involved at least one parent with a high *gca* (high × high, high × low) for the trait in question. The current results are in line with those reported elsewhere [36] based on watermelon crosses with high *gca* effects from parents with either high × high or high × low *gca* effects. The crossover having a high significant value of *sca* involving two best general combiners, would be due to the additive gene action and the additive × non-additive gene interaction which are fixable [30, 37]. For a given character, the best combination in terms of *gca* obtained with a cross involving a single parent having a high *gca* is the result of the additive gene interaction × dominance in the expression of the trait [38] and therefore, the diversity of values of parental *gca* would play an important role in the production of hybrids with significant *gca* values [30, 39]. However, for a given trait, parents with poor general combining skills sometimes give good combinations when crossed with each other, as is the case here in the cross between *Zouan-H × Long* for the weight of the fruit and the weight of the pulp. Significant *sca* effects resulting from crosses of parents with low *gca* indicate the presence of epistasis (non-allelic interaction) at heterozygous loci that are not fixable [38]. It is therefore suggested to use these crosses for the selection of a plant in later generations of breeding [38]. It should also be noted that non-significant *sca* can be obtained after a crossover involving two best general combiners [8]. Therefore, it cannot be generalized that parents with high *gca* effects could only produce good hybrids [40]. Furthermore, combinations having recorded high sca values can be improved through conventional genetic improvement methods such as biparental crosses and selective diallel crosses, subsequently, followed by the pedigree selection method, to break any epistatic link that may occur, in order to isolate transgressive segregants [41, 42, 39].

## 5. Conclusion

Our study helped to test four parental lines of *Cucurbita moschata* D. and their 12 F1 hybrids from a complete diallel cross for the evaluation of fruit and seed traits and plant height. It appears from our work that there is very high genetic diversity among the parental lines *Soubre, Long, Oval*, and *Zouan-H* and their descendants. Heterosis effects relative to the average parent were observed in hybrids for all traits except fruit length. With respect to the best parent, heterosis for different combinations of parents was significant for all traits except plant height, number of fruit per plant and fruit length. Significant *gca* effects were observed for all traits for different hybrid combinations. The traits studied were predominantly under the influence of genes with additive effects. The *Soubre* parental line proved to be the best combiner for several traits. However, *Zouan-H* was the best combiner for the number of seeds while *Long* was the best combiner for vegetative development and *Oval* was the best candidate for the mass of fresh seeds. Significant sca effects were recorded at some crosses for each trait, suggesting that these crosses can be used for the development of new and more productive hybrids.

## Notes

### Competing Interest Statement

The authors have declared no competing interest.

## References

1. FAO (2013). Cereal Supply and Demand Brief (http://www.fao.org/worldfoodsituation /wfs-home/csdb/en/).

2. Megueni C, Awono ET, and Ndouenkeu R (2011). Effet simultané de la dilution et de la combinaison du Rhizobium et des mycorhizes sur la production foliaire et les propriétés physico-chimiques des jeunes feuilles de Vigna unguiculata (L.) J. Appl. Biosci. 40 (1): 2668–2676.

3. Zoro BIA, Koffi KK, and Djè Y (2003). Botanical and agronomic characterization of three species of cucurbites consumed in West African sauce: Citrullus sp., Cucumeropsis mannii Naudin and Lagenaria siceraria (Molina) Standl. Biotechnol. Agron. Soc. Environ. 7(3–4), 189–199.

4. Abdein MAE, Hassan HMF and Dalia HM (2017). General performance, combining abilities and yield component traits in Pumpkin (Cucurbita moschata Poir.) at different conditions. Current applied science and technology. 17 (1): 121–129.

5. Jahan TA, Islam A, Rasul MG, Mian MAK, and Haque MM (2012). Heterosis of Qualitative and Quantitative Characters in Sweet Gourd (Cucurbita Moschata Duch.Ex Poir). African Journal of Food, Agriculture, Nutrition and Development 12(3): 6186–6199.

6. Jahan M, Koocheki A, Tahami MK, Amiri MB, and Nassiri MM (2012). The effects of simultaneous application of different organic and biological fertilizers on quantitative and qualitive characteristics of Cucurbita pepo L. Journal of Life Sciences 12 (1): 1145–1149.

7. Enneb S, Drine S, Bagues M, Triki T, Boussora F, Guasmi F, Nagaz K, and Ferchichi A (2020). Phytochemical profiles and nutritional composition of squash (Cucurbita moschata D.) from Tunisia. South African Journal of Botany 130: 165–71.

8. Brou KF, Sie RS, Adjoumani K, Saraka YDM, Kouago BA and Adahi CD. (2018). Diallel analysis in Citrullus mucosospermus (Fursa) for fruit traits. African Journal of Agricultural Research 13 (38): 2006–2015.

9. Griffing B (1956). Concept of general and specific combining ability in relation to diallel crossing systems. Australian Journal of Biological Sciences 9(4): 463–493.

10. Feyzian E, Dehghani H, Rezai AM and Javaran MJ (2009). Diallel cross analysis for maturity and yield-related traits in melon (Cucumis melo L.). Euphytica 168:215–223. DOI 10.1007/s10681-009-9904-9.

11. Acquaah G (2007). Principles of Plant Genetics and Breeding. Oxford:Wiley-Blackwell.

12. Viana JMS and Matta FP (2003). Analysis of general and specific combining abilities of popcorn populations, including selfed parents. Genet. Mol. Biol. 26:465–471. doi:10.1590/S1415-475720030004 00010.

13. Yu K, Wang H, Liu X, Xu C, Li Z, Xu X, Liu J, Wang Z and Xu Y (2020). Large-Scale Analysis of Combining Ability and Heterosis for Development of Hybrid Maize Breeding Strategies Using Diverse Germplasm Resources. Front. Plant Sci. 11:660. doi: 10.3389/fpls.2020.00660.

14. Backer RJ (1978). Issues in diallel analysis. Crop Sci. 18: 533–536.

15. Hung HY and Holland JB (2012). Diallel analysis of resistance to Fusarium ear rot and fumonisin contamination in maize. Crop Sci. 52:2173–2181. doi:10.2135/cropsci2012.03.0154.

16. Wynne JC, Emery DA and Rice Pw (1970). Combining ability estimates in Arachis hypogaea L. II. Field performance of F1 hybrids. Crop Sci. 10 (1): 713–715.

17. Heydari F, Ramezanpour SS, Soltanloo H, Mehdi KA and Shaban K (2012). Estimation of genetic parameters and effective factors in resistance to septoria leaf blotch of wheat under field condition. International Journal of Agriculture and Crop Sciences 4(18): 1376–1384.

18. Smith HH (1952). Fixing transgressive vigor in Nicotiana rustica. In: Goodwin TW ed. Heterosis. Ames, IA, Iowa State College Press. Pp. 161–174.

19. Zeinanloo A, Shahsavari A, Mohammadi A and Naghavi MR (2009). Variance component and heritability of some fruit characters in olive (Olea europaea L.). Scientia Horticulturae 123:68–72.

20. R Core Team (2023). _R: A Language and Environment for Statistical Computing_. R Foundation for Statistical Computing, Vienna, Austria. <https://www.R-project.org/.

21. Onofri A, Terzaroli N and Russi L (2020). Linear models for diallel crosses: a review with R functions. Theor. Appl. Genet. 10.1007/s00122-020-03716-8.

22. Paradis E and Schliep K (2019). “ape 5.0: an environment for modern phylogenetics and evolutionary analyses in R.” _Bioinformatics_, *35*, 526–528. doi:10.1093/bioinformatics/bty633 <10.1093/bioinformatics/bty633>.

23. Letunic I and Bork P (2021). Interactive Tree Of Life (iTOL) v5: an online tool for phylogenetic tree display and annotation. Nucleic Acids Res. 2:49(W1):W293–W296. doi: 10.1093/nar/gkab301. PMID: 33885785; PMCID: PMC8265157.

24. Ananda G (2015). Courges et Courgettes. Légumes et Fruits, Kokopelli 1(1) :1–17.

25. Seka D, Kouago BA, Bonny BS (2023). Assessment of the variability of the morphological traits and differentiation of Cucurbita moschata in Cote d’Ivoire. Sci Rep. 2023 Mar 6;13(1):3689. doi: 10.1038/s41598-023-30295-7. PMID: 36878922; PMCID: PMC9988981.

26. Ahsan, M.Z., Majidano, M.S., Bhutto, H., Soomro, A.W., Panhwar, F.H., Channa, A.R., and Sial, K.B. (2015). Genetic variability, coefficient of variance, heritability and genetic advance of some Gossypium hirsutum L. accessions. J. Ag. Sci. 7 (2):147– 151.

27. Bello OB, Aminu D, Gambo A, Azeez AH, Lawal M, Ali I, and Abdulhamid UA (2015). Genetic diversity, heritability and genetic advance in okra [Abelmoschus esculentus (L.) Moench]. Bangladesh Journal of Plant Breeding and Genetics, 28 (2):25–38.

28. Golabadi M, Golkar P and Eghtedary A (2015). Combining ability analysis of fruit yield and morphological traits in greenhouse cucumber (Cucumis sativus L.). Canadian Journal of Plant Science. 95 (2):377–385.

29. Adjoumani K, Kouonon LC, Koffi GK, Bony BS, Brou KF, Akaffou DS and Sié RS. (2016) Analysis on genetic variability and heritability of fruit characters in Citrullus lanatus (Thunb.) Matsumura and Nakai (Cucurbitaceae) cultivars. J. Anim. Plant Sci 28 (1): 4340–4355.

30. Banerjee PP, and Kole PC (2009). Analysis of genetic architecture physiological characters in sesame (Sesamum indicum L.). Euphytica, 168:11–22.

31. Yousfi K (2011). Etude agronomique et analyse diallèle de six variétés de blé dur (Triticum durum Desf.). Mémoire soutenu le 17/11/2011 à l’Ecole Nationale Agronomique El-Harrach-Alger en vue de l’obtention du diplôme magistère. 65 pages.

32. Mohammadi AA, Saeidi G and Arzani A (2010). Genetic analysis of some agronomic traits in flax (Linum usitatissimum L) Australian Journal of Crop Sci. 4(5):343–352.

33. Wannows AA, Sabbouh MY and Al-Ahmad SA (2015). Generation mean analysis technique for determining genetic parameters for some quantitative traits in two maize hybrids (Zea mays L.). Jordan Journal of Agricultural Sciences 11 (1): 59–73.

34. Thingamajig C and Pugalendhi L (2013). Heterosis studies in Bitter Gourd for yield and related. International Journal of Vegetable Science, 19 (2): 109–125.

35. Masny A, Madry W and Żurawicz E. (2005). Combining ability analysis of fruit yield and fruit quality in ever-bearing strawberry cultivars using an incomplete diallel cross design. Journal of Fruit and Ornamental Plant Research. 13(1): 5–17.

36. Hatem AK (2009). Heterosis and combining ability for yield and some characters in watermelon (Citrullus lanatus, Thunb). Minfya J. Agric. Res., 34 (1): 147–162.

37. Adarsh MN and Kumari P (2015). Combining ability and gene action studies for important horticultural traits in chilli (Capsicum annuum L.). International Journal of farm Sciences 5(1): 251–262.

38. Nataša L, Sofija P, Miodrag D, Hristov N, Vukosavljev M and Srećkov Z (2014). Diallel analysis for spike length in winter wheat. Turkish Journal of Agricultural and Natural Sciences 2 (1): 1455–1459.

39. Venel C (2022). Analyse du déterminisme génétique de l’aptitude à la combinaison et de l’effet des hétérosis des caractères liés au rendement chez labetterave sucrière en utilisant les designs North Carolina II et North Carolina III. Sciences du Vivant (q-bio), dumass-03855426 :1–100.

40. El-Tahawey MAFA, Kandeel AM., Youssef SMS and Abd El-Salam MMM (2015). Heterosis, potence ratio, combining ability and correlation of some economic traits in diallel crosses of pumpkins. Egypt. J. Plant Breed. 19(2):419 – 439 (2015).

41. Iqbal AM, Nehvi FA, Wani SA, Dar ZA, Lone AA and Qadri H (2015). Combining ability study over environments in dry beans (Phaseolus vulgaris L.) SAARC Journal of Agriculture 10(2): 61–69.

42. Fellahi Z, Hannachi A, Oulmi A and Bouzerzour H (2018). Analyse des aptitudes générale et spécifique à la combinaison chez le blé tendre (Triticum aestivum L.). Revue Agriculture. 9(1):60 –70.

